# Automated phenol-chloroform extraction of high molecular weight genomic DNA for use in long-read single-molecule sequencing

**DOI:** 10.1101/2022.01.26.477939

**Authors:** Andrew W. Liu, Alejandro Villar-Briones, Nicholas M. Luscombe, Charles Plessy

**Affiliations:** Genomics & Regulatory Systems Unit, Okinawa Institute of Science & Technology Graduate University, 1919-1 Tancha, Kunigami-gun, Okinawa Japan 904-0425; Instrument Analysis Section, Okinawa Institute of Science & Technology Graduate University, 1919-1 Tancha, Kunigami-gun, Okinawa Japan 904-0425

**Keywords:** Automation, Robotics, Organic Extraction, DNA extraction, Genome Sequencing

## Abstract

In order to automate the genome sequencing pipeline in our laboratory, we programmed a dual-arm anthropomorphic robot, the Robotic Biology Institute’s Maholo LabDroid, to perform organic solvent-based genomic DNA extraction from cell lysates. To the best of our knowledge, this is the first time that automation of phenol-chloroform extraction has been reported. We achieved routine extraction of high molecular weight genomic DNA (>100 kb) from diverse biological samples including algae cultured in sea water, bacteria, whole insects, and human cell lines. The results of pulse-field electrophoresis size analysis and the N50 sequencing metrics of reads obtained from Nanopore MinION runs verified the presence of intact DNA suitable for direct sequencing. We present the workflow that can be used to program similar robots and discuss the problems and solutions we encountered in developing the workflow. The protocol can be adapted to analogous methods such as RNA extraction, and there is ongoing work to incorporate further post-extraction steps such as library construction. This work shows the potential for automated robotic workflows to free molecular biological researchers from manual interventions in routine experimental work. A time-lapse movie of the entire automated run can be seen at http://doi.org/10.6084/m9.figshare.17097065

## Background

DNA extraction is a routine, yet critical step in genomics research. Across diverse settings and applications, the purity of isolated DNA affects the sensitivity of downstream processes such as PCR and sequencing. In addition, recent developments in sequencing technologies have increased the emphasis of extracting intact or high-molecular weight genomic DNA from a wide variety of biological materials. The Oxford Nanopore Technologies (Kasianowicz *et al*., 1996; Jain *et al*., 2018) and PacBio Single Molecule Real-Time (Eid *et al*., 2009; Chin *et al*., 2013) platforms enable users to obtain long sequence reads – a major limiting factor being the availability of long DNA molecules. Sequencing data from such platforms are routinely used to generate whole genome sequence assemblies.

Organic, or phenol-chloroform, extraction is one of the oldest and, for many years, the most widely used method for DNA isolation. A phenol-chloroform mixture is added to lysed or homogenised biological material. When centrifuged, the unwanted proteins and cellular debris are separated away in the organic phase, and DNA molecules in the aqueous phase can be cleanly isolated for analysis. Among the many applications, organic extraction was historically used to isolate nucleic acids from viruses, which had proved challenging owing to the chemically resistant protein coats surrounding their genomes (Sinsheimer, 1959; Thomas & Berns, 1961; Saito & Miura, 1963). Despite the development of modern extraction approaches, the phenol-chloroform method continues to be relevant because it works reliably for many biological samples and consistently gives high yields of high molecular weight DNA (Ghaheri *et al*. 2016; Bouso & Planet, 2019; Torii *et al*. 2021). However, it is also time-consuming, involves the use of hazardous organic solvents, and requires samples to be transferred between multiple tubes, increasing the risk of error or contamination.

Here, to establish consistent isolation of high molecular weight genomic DNA and reduce the amount of manual work needed for a routine protocol involving toxic reagents, we programmed the Robotic Biology Institute’s Maholo LabDroid, a dual-arm anthropomorphic robot, to perform organic genomic DNA extraction from lysed cells and homogenized tissues.

The LabDroid is an example of a versatile modular system that has demonstrated a practical application in a laboratory environment (Ochiai *et al*., 2020). Maholo’s design allows users to design a workspace/equipment layout within the reach of the robot (Figure 1). Automating the procedure was expected to enhance reproducibility, increase throughput, and reduce the possibility of DNA shearing into shorter fragments by minimizing excessive handling. Devices for automated DNA isolation are available, but to the best of our knowledge, this is the first report that a phenol-chloroform extraction has been automated.

**Figure 1.**
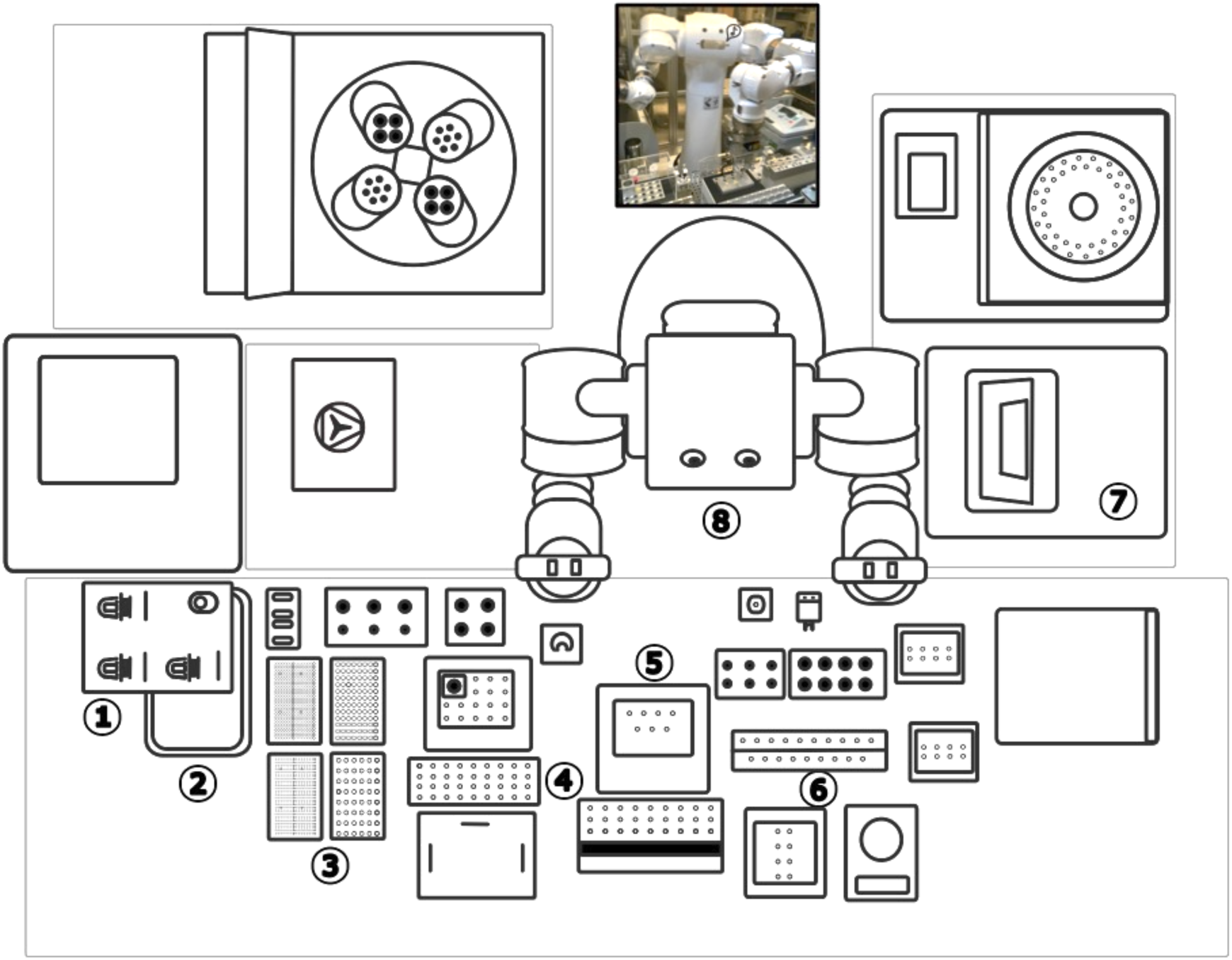
Schematic overhead view of the LabDroid’s workspace layout. 1) micro-pipettor station 2) waste pipette tip pan 3) pipette tips 4) reagents/tube rack 5) 4°C cool block 6) sample tube rack/mixing station 7) high speed centrifuge 8) robot.

## Material & Methods

### Ethical considerations

The human cell lines, laboratory animals and microbial stocks used in this study do not require ethics approval in accordance with Okinawa Institute of Science & Technology Graduate University (OIST)’s Animal Care and Use Committee and Japan Society for the Promotion of Science (JSPS) guidelines. All procedures regarding the welfare and handling of organisms meet ARRIVE guidelines for the care and use of experimental animals. In particular, Drosophila were anaesthetized with carbon dioxide (CO_2_) prior to mechanical cell disruption.

### Sample preparation

Samples were manually homogenized either by grinding using a simple tube-based mortar and pestle (single whole eclosed fruit fly) or freeze-thawing 5 times (cultured cyanobacteria and algae). The human cell culture was not subjected to mechanical disruption.

Newly eclosed single *Drosophila melanogaster* were isolated in vials containing food but no yeast for 18 hours, and then ground gently in 200 µl ice-cold extraction buffer (50mM Tris-Cl pH 8.0, 100 mM NaCl, 2.5 mM EDTA, 0.5% SDS, 0.2 mg/ml Proteinase-K, 1% 2-mercaptoethanol) with a tube-based mortar & pestle on ice. The homogenized extract was incubated overnight for 18 hours at 55°C. The cell lysate was centrifuged at 11,000g (Tomy, MX-302) for 10 minutes at 4°C and 200 µl of the cleared lysate used for robotic extraction.

Cultured human embryonic kidney cells, HEK293, were dissociated from a confluent tissue culture flask with 0.25% Trypsin-EDTA for 3 minutes at 37°C (Gibco, 25200056), counted on a hemocytometer, centrifuged at 800g for 5 minutes and washed with 10 ml PBS. 1 × 10^6^ cells were drawn off and spun again at 800g for 5 minutes. The PBS was aspirated, and the cell pellet was resuspended in extraction buffer 50 mM Tris-Cl pH 8.0, 100 mM NaCl, 10 mM EDTA, 0.5% SDS, 0.2 mg/ml Proteinase-K, 1% 2-mercaptoethanol and incubated for 2 hours at 55°C. The cell lysate was processed immediately by centrifugation at 11,000g for 10 minutes at 4°C and 200 µl of the cleared lysate used for robotic extraction.

The cyanobacterium *Synechoccous sp*. was centrifuged at 2,500g for 5 minutes from a one-month-old culture; the resulting pellet was washed with filtered autoclaved seawater three times and gently resuspended in 250 µl of extraction buffer (50 mM Tris-Cl pH 8.0, 100 mM NaCl, 20 mM EDTA, 1% SDS, 2% 2-mercaptoethanol) and frozen overnight at –80°C. The next morning the cyanobacteria were thawed on ice, an equal volume of 5%SDS, 50 mM EDTA & 50 mM Tris-Cl pH 8 was mixed by inversion and subjected to four more freeze/thaw cycles. Mixed algal cultures of 6×10^6^ *Chaetoceros calcitrans*, 1 × 10^7^ *Isochrysis sp*. and 1 × 10^6^ *Rhinomonas reticulata* were centrifuged similarly from a one-month-old culture; the resulting pellet washed with filtered autoclaved seawater three times and gently resuspended in 250 µl of ice-cold extraction buffer (100 mM Tris-Cl pH 8.0, 1.4 M NaCl, 20 mM EDTA, 2% cetyltrimethylammonium bromide, 0.1% polyvinylpolypyrrolidone, 2% 2-mercaptoethanol) and then frozen overnight at – 80°C. The next morning, the algal extractions were thawed on ice and subjected to four more freeze/thaw cycles. All extractions were centrifuged at 4°C for 10 min at 15,000g and 200 µl of the cleared lysate used for robotic extraction.

### Robot-assisted organic extraction

A workflow based on a modified organic solvent based back-extraction method (Giles *et al*., 1980) was programmed into the LabDroid with Robotic Biology Institute’s BioPortal ProtocolMaker graphical user interface (<u>10.5281/zenodo.5733820</u>). Though the robot is capable of processing 14 samples in a run, the development was carried out with three samples. The procedural schematic and robotic workflow are outlined in Figure 2A and 2B, respectively. To start, 200 µl of cleared lysate was transferred to a new 2 ml tube (Eppendorf 0030 120.094) and used for the robot-assisted protocol. 200 µl of Phenol, chloroform and isoamyl alcohol (25:24:1, Sigma-Aldrich P3803) was used for the first two robot “hand-mixed” organic extractions. The closed-capped tubes were mixed by the robot’s arm/manipulators by inversion at a rate of one every 3 seconds, for a total of 5 minutes. These were later centrifuged at 15,000g for 1 minute at 4°C. The first extraction aqueous phase was transferred to a new 2 ml Eppendorf tube awaiting at 4°C. Remaining DNA left behind was back-extracted by the same means after addition of fresh extraction buffer to the phenol phase and the second aqueous phase was pooled in same 2 ml tube with the first aqueous phase. 400 µl of chloroform (Wako 3034-02603) was saturated with 10 mM Tris-Cl pH 8.5 1 mM EDTA and then used to extract the pooled aqueous phases. These were “hand-mixed” and centrifuged similarly as the phenol extractions. The 325 µl of the aqueous phase from the chloroform extraction was then transferred to a new 1.5 ml centrifuge tube (Eppendorf 0030 125.215). 650 µl of 100% ethanol and 33 µl of 3 M sodium acetate pH 5.2 (Sigma Aldrich, S7899) was added, “hand-mixed” for 5 minutes and centrifuged at 15,000g for 30 minutes at 4°C. 50 µg glycogen (Thermo, R0561) was added to the final 1.5 ml tubes before the start of the robotic run to help visualize pellets after precipitation and centrifugation. After centrifugation, the tubes were placed into a 4°C block awaiting sample recovery. An entire run was video-captured, and the steps are highlighted at key points described in this manuscript as a visual demonstration of the robot’s movement (<u>10.6084/m9.figshare.17097065</u>)

**Figure 2.**
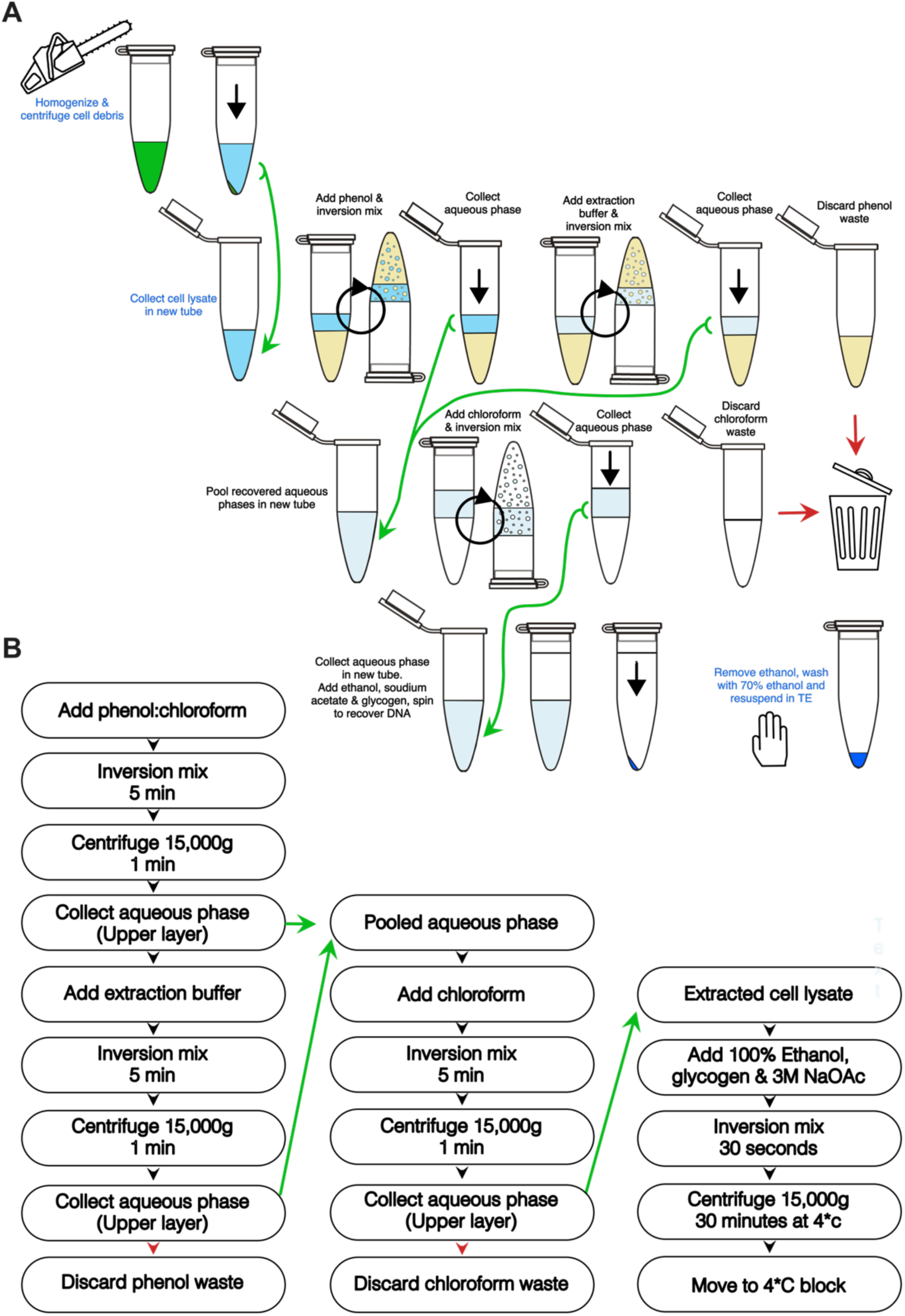
Graphical description of the LabDroid’s workflow. **A**, Schematic representation of manual (blue text) and robotic (black text) manipulations. Circular black arrows indicate “hand-mixing” inversions and black arrows, centrifugation. **B**, Automated workflow of extraction protocol. For both panels, green arrows indicate transfer to new tube and red arrow heads, disposal of waste.

### Pulsed field electrophoresis and long-read sequencing

Subsequently, the ethanol supernatant from the last automated step was removed from the pelleted DNA by manual aspiration. The pellets were carefully washed with 70% ethanol, removed by manual aspiration and air dried for 5 minutes at the bench. The washed DNA pellets were gently resuspended in 15 µl 10 mM Tris-Cl and 1 mM EDTA and quantitated with Qubit high-sensitivity double-stranded DNA reagents and fluorometer (Thermo Fisher, Q32851). 10 ng of the sample was used for pulsed-field (Agilent, FP-1002-0275) or standard capillary electrophoresis Tape Station (Agilent, 5067-5365). 1000 ng of gDNA was used for adapter ligation (Oxford Nanopore Technologies, ONT LSK-109). Between 40–500 ng of adapter ligated DNA was loaded onto a MinION R9.4 flow cell (ONT FLO-MIN106D) and allowed to run for 18-24 hours. N50 metrics were determined with the MinKNOW software (ONT).

## Results

Movie 1 shows an entire run was of LabDroid’s extraction process, and the steps are highlighted at key points described in this manuscript as a visual demonstration of the robot’s movement (<u>10.6084/m9.figshare.17097065)</u>.

### Length of genomic DNA from automated preparations

Mammalian, algal, cyanobacteria and drosophila genomic DNAs were prepared using the organic extraction protocol programmed for Maholo. Long-read sequence data were confirmed after analysis of completed Nanopore MinION using the standard sequencing N50 metric for the length of average read: N50 values were 30-40 kb for algal samples, 20kb for cyanobacteria and 16kb for drosophila (Table 1). Representative results from the reverse-field gel electrophoresis genomic analysis kits are shown for *Synechecoccus* and *Rhinomonas* preparations (Figure 3A & 3B, respectively). For all preparations, fragments ranged in size between 50kb and 200kb, similar to manually prepared DNA in the hands of an experienced technician.

**Table 1.** Representative Nanopore MinION N50 raw-reads and fragment length analysis

| Organism | Sample Material | N50 (kb) | Peak Fragment Size (kb) |
| --- | --- | --- | --- |
| *Synechococcus sp.* | Bacterial culture | 19.13 | 152.6 |
| *Chaetoceros calcitrans* | Algal culture | 35.55 | >165 |
| *Isochrysis sp.* | Algal culture | 39.67 | >165 |
| *Rhinomonas reticulata* | Algal culture | 29.54 | 152.2 |
| *Drosophila melanogaster* | Whole flies | 16.43 | >60 (Tape Station) |
| *Homo sapiens (HEK293)* | Tissue culture | ND | 120.5 |

**Figure 3.**
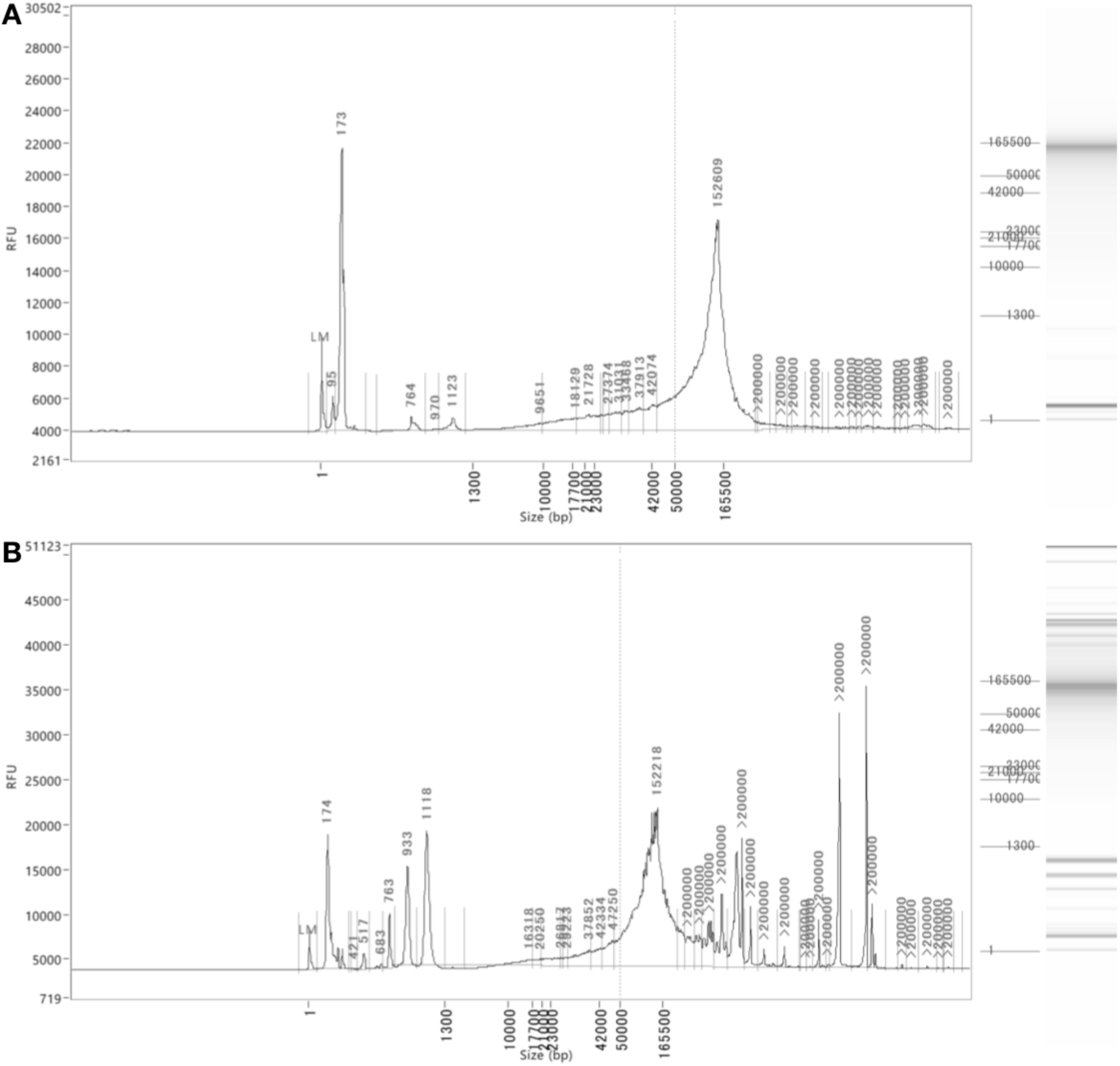
Representative Femto Pulse fragment length results from: **A**, cyanobacteria (*Synechococcus sp*.) and **B**, microalgae (*Rhinomonas reticulata*).

### Troubleshooting problems during protocol development

We solved practical problems iteratively in order to implement a standard phenol-chloroform DNA extraction method on the Maholo LabDroid. Below we describe the major obstacles we encountered and their solutions (Table 2).

**Table 2.**
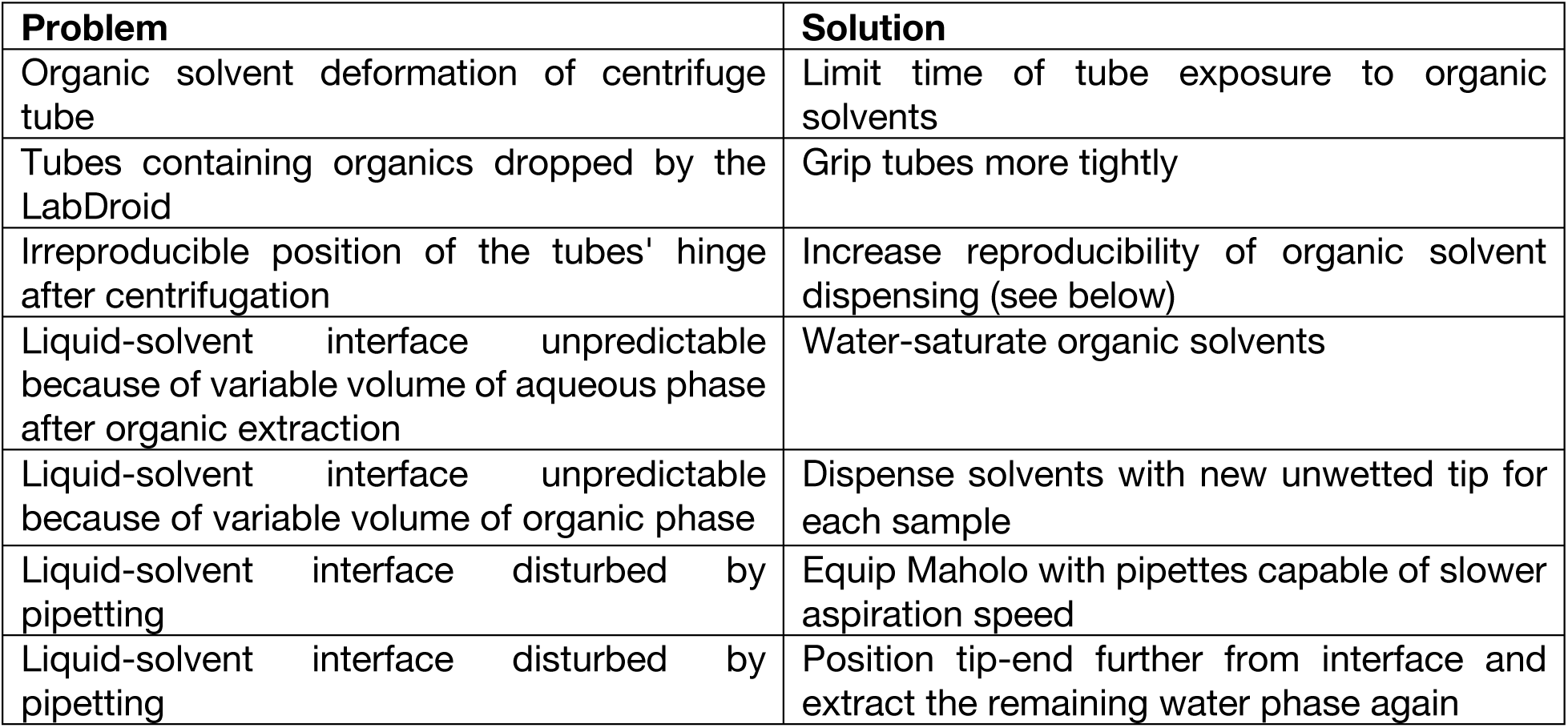
Summary of problems encountered and solutions.

Organic solvents can affect the physical properties of plastics even though the effects may be imperceptible to the naked eye. Here, we found that prolonged exposure of Eppendorf tubes to phenol and chloroform caused enough changes in their dimensions for the LabDroid’s manipulators to sense and abort the run. We overcame this issue by adding aliquoted phenol and chloroform just before the start of the organic extractions. Furthermore, we found that the manipulators would drop the tubes owing to the changes in tube elasticities after centrifuging with organic solvent. The force to grip the tubes was increased during tube removal from the centrifuge to resolve this problem.

As the LabDroid does not detect the position of the liquid interfaces, it relies on the preprogrammed volumes to be correct. Mixture of organic solvents and water can vary in final volume depending on how much water was originally present, for instance through contamination of solvent aliquots by the moisture in the air. Variations in volumes reduce the precision of the robot’s pipetting capabilities relative to the phase interface. It also impaired the robot’s handling of the centrifugation step since the slight imbalance caused individual tubes to rotate about their vertical axes as they balanced the mass distribution in the rotor. This changed the position of the hinge at the end of the run which prevented the robot from removing the rotated tube from the rotor. To ensure volume reproducibility, chloroform was 10 mM Tris-Cl pH 7.5 1 mM EDTA (TE) saturated the night before (1:1 phenol and chloroform was purchased pre-equilibrated with TE pH 8) to prevent any aqueous phase volume loss due to the solvent’s slight water solubility. Finally, we also programmed the LabDroid to change the pipette tips after each aliquot of solvent was added, as the actual volume of solvent transferred by the tips was slightly greater after the tip had been pre-wetted.

The greatest hurdle was the collection of aqueous phases after extraction and centrifugal phase separation; this occurs three times in the protocol. The LabDroid initially was fitted with a brand of Bluetooth actuated liquid handling micro-pipettors that could not aspirate the aqueous phase without disturbing the organic/aqueous interface. As a result, the aqueous phases were contaminated with solvent which caused purity problems downstream. This was solved when new electronic pipettors (Viaflow, Integra Biosciences), capable of slower rates of aspiration, were outfitted to replace the earlier model. The aspiration offset height (the distance between the end of the pipette’s tip to the bottom of the tube) was also increased by 1 mm to avoid disturbing the organic/aqueous interface. As this increased the dead volume, we added a back-extraction step to collect the leftover aqueous phase.

### Attempts at extending the protocol beyond organic extraction steps

Currently, the automated protocol ends with the pelleted DNA in salted ethanol solution, which is suitable for long-term DNA storage. We tried to incorporate a last step to precipitate the pellet in 70% ethanol and resuspend it in TE. We were unsuccessful as the pellet dislodged after the wash centrifugation. While a skilled technician can carefully remove supernatant in the presence of a floating pellet, a robot with no visual sensors cannot. To overcome this, we tried partial aspiration of the 70% ethanol wash and drying 50-70 µl of supernatant with a vacuum centrifuge for 40 minutes at room temperature at 0.02 bar, but we found this method partially degraded preparations. As a result, we decided to leave out the 70% ethanol wash from the automated protocol.

## Discussion

Our main challenges in developing the automated protocol were problems that an experienced human investigator tends to resolve instinctively, but which prevents a deterministically programmed robot from continuing to the next action, thus aborting the run.

We did not include the tissue homogenization and digestive steps in the automation because they involve long incubation times during which the robot would be idle. More notably, different tissue samples and organisms require different treatments to free the DNA from cells. Nevertheless, it is conceivable to use the platform to determine the optimal incubation for groups of samples of equivalent properties, in the same spirit as the work of Kanda *et al*., (2020) which reported AI-based optimization of iPS cell-culture conditions with Maholo. Further improvements of the method, such as the use of wide-bore tips to reduce DNA shearing, require more extensive reprogramming of the LabDroid’s movements as it currently relies on a fixed shape for all its tools and consumables.

The LabDroid’s gentle mixing of the organic extractions supplied the ideal amount of agitation necessary to create an organic-aqueous emulsion that resulted in a clean long-read DNA preparation. The results obtained have given us confidence in obtaining intact genomic DNA for routine genome sequencing and assembly, and we have begun using the robot in our sample preparation-to-sequence analysis workflow thus shortening benchwork and reducing exposure to organic solvents of the researcher.

With only minor changes, such as preparative equilibration of the phenol component to pH 4.5 with a small volume of 3M sodium acetate, the protocol can be adapted to total RNA with no further modifications to the robotic workflow (Wallace, 1987). With further development, we hope to program the LabDroid to perform the entire workflow from extraction to loading the adapter-ligated DNA on Nanopore MinION flow-cells.

<u>10.6084/m9.figshare.17097065</u>

**Movie 1**. Video showing the LabDroid in action through the automated DNA extraction protocol

## Underlying Data Links

Job Files for LabDroid operations: <u>10.5281/zenodo.5733820</u>

Movie of LabDroid run: <u>10.6084/m9.figshare.17097065</u>

ONT Nanopore MinKNOW Metrics and Agilent fragment size analysis: <u>10.5281/zenodo.5921205</u>

## Acknowledgments

We would like to thank Dr. Takakazu Yokokura (Formation and Regulation of Neuronal Connectivity Research Unit, Okinawa Institute of Science and Technology) for supplying flies, Dr. Narae Kim (Nucleic Acid Chemistry and Engineering Unit, OIST) for providing HEK293 cells, Mayumi Kawamitsu (Sequencing Section, OIST) for providing Femto Pulse results, and Drs. Motohisa Kamei & Shigeji Tasaka (RBI) for input during manuscript drafts.

## References

1. Complete nontuberculous mycobacteria whole genomes using an optimized DNA extraction protocol for long-read sequencing. Bouso, J.M., Planet, P.J. BMC Genom. 20, 793 (2019). DOI: 10.1186/s12864-019-6134-y

2. Nonhybrid, finished microbial genome assemblies from long-read SMRT sequencing data. Chin C.S., Alexander D.H., Marks P., Klammer A.A., Drake J., Heiner C., Clum A., Copeland A., Huddleston J., Eichler E.E., Turner S.W., Korlach J. Nat Methods. 2013 Jun;10(6):563–9. DOI: 10.1038/nmeth.2474 Epub 2013 May 5. PMID:23644548.

3. Real-Time DNA Sequencing from Single Polymerase Molecules. Eid, J., Fehr, A., Gray, J., Luong, K., Lyle, J., Otto, G., Peluso, P., Rank, D., Baybayan, P., Bettman, B., Bibillo, A., Bjornson, K., Chaudhuri, B., Christians, F., Cicero, R., Clark, S., Dalal, R., deWinter, A., Dixon, J., Foquet, M., Gaertner, A., Hardenbol, P., Heiner, C., Hester, K., Holden, D., Kearns, G., Kong, X., Kuse, R., Lacroix, Y., Lin, S., Lundquist, P., Ma, C., Marks, P., Maxham, M., Murphy, D., Park, I., Pham, T., Phillips, M., Roy, J., Sebra, R., Shen, G., Sorenson, J., Tomaney, A., Travers, K., Trulson, M., Vieceli, J., Wegener, J., Wu, D., Yang, A., Zaccarin, D., Zhao, P., Zhong, F., Korlach, J., & Turner, S. Science 2009, 323, 133–138. DOI: 10.1126/science.1162986

4. A comparative evaluation of four DNA extraction protocols from whole blood sample. Ghaheri M., Kahrizi D., Yari K., Babaie A., Suthar R.S., Kazemi E. 2016; Cell. Mol. Biol. 62(3): 120–124 PMID: 27064884

5. Maternal inheritance of human mitochondrial DNA. Giles R.E., Blanc H., Cann H.M., Wallace D.C. Proc. Natl. Acad. Sci. U.S.A. 1980, 77 (11): 6715–6719; DOI: 10.1073/pnas.77.11.6715

6. Nanopore sequencing and assembly of a human genome with ultra-long reads. Jain M., Koren S., Miga K.H., Quick J., Rand A.C., Sasani T.A., Tyson J.R., Beggs A.D., Dilthey A.T., Fiddes I.T., Malla S., Marriott H., Nieto T., O’Grady J., Olsen H.E., Pedersen B.S., Rhie A., Richardson H., Quinlan A.R., Snutch T.P., Tee L., Paten B., Phillippy A.M., Simpson J.T., Loman N.J., Loose M. Nat Biotechnol. 2018 Apr;36(4):338–345. DOI: 10.1038/nbt.4060 Epub 2018 Jan 29. PMID: 29431738; PMCID: PMC5889714

7. Robotic Search for Optimal Cell Culture in Regenerative Medicine. Kanda G.N., Tsuzuki T., Terada M., Sakai N., Motozawa N., Masuda T., Nishida M., Watanabe C.T., Higashi T., Horiguchi S.A., Kudo T., Kamei M., Sunagawa G.A., Matsukuma K., Sakurada T., Ozawa Y., Takahashi M., Takahashi K., Natsume T. bioRxiv 2020.11.25.392936; DOI: 10.1101/2020.11.25.392936

8. Characterization of individual polynucleotide molecules using a membrane channel. Kasianowicz J.J, Brandin E., Branton D., Deamer D.W. Proc. Natl. Acad. Sci. U.S.A. 1996; 93 (24): 13770–13773; DOI: 10.1073/pnas.93.24.13770

9. A variable scheduling maintenance culture platform for mammalian cells. Ochiai K., Motozawa N., Terada M., Horinouchi T., Masuda T., Kudo T., Kamei M., Tsujikawa A., Matsukuma K., Natsume T., Kanda G.N., Takahashi M., Takahashi K. SLAS (Society for Laboratory Automation and Screening) Technol. 2021; 26(2):209–217 DOI: 10.1177/2472630320972109

10. Preparation of transforming deoxyribonucleic acid by phenol treatment. Saito H., Miura K. Biochim. Biophys. Acta 1963; 72: 619–629 DOI: 10.1016/0926-6550(63)90386-4

11. A single-stranded deoxyribonucleic acid from bacteriophage φX174. Sinsheimer R.L. J. Mol. Biol. 1959; 1(1): 43–53 DOI: 10.1016/S0022-2836(59)80006-1

12. The physical characterization of DNA molecules released from T2 and T4 bacteriophage. Thomas C.A. & Berns K.I. J. Mol. Biol. 1961; 3(3): 277–288. DOI: 10.1016/S0022-2836(61)80069-7

13. Applicability of polyethylene glycol precipitation followed by acid guanidinium thiocyanate-phenol-chloroform extraction for the detection of SARS-CoV-2 RNA from municipal wastewater. Torii S., Furumai H., Katayama H. Sci. Total Environ. 2021 Feb; 756:143067. DOI: 10.1016/j.scitotenv.2020.143067

14. Large and small phenol extractions. Wallace DM; Methods in Enzymology 1987, 152:33–41 DOI: 10.1016/0076-6879(87)52007-9

